# Selection of horizontal gene transfer through public good production

**DOI:** 10.1101/315960

**Authors:** Tatiana Dimitriu, Dusan Misevic, Julien Benard Capelle, Ariel B Lindner, Sam P Brown, Francois Taddei

## Abstract

In bacteria, cooperative genes encoding public good molecules are preferentially located on mobile genetic elements (MGEs), and horizontal transfer of MGEs favours the maintenance of public good cooperation. The rate of horizontal transfer itself can evolve in response to selective pressures acting on both MGEs and bacterial hosts: benefits and costs of infectious spread, but also indirect effects of MGE genes to the host. We show here that carriage of public good genes on MGEs can generate another indirect selection for MGE transfer. Transfer increases public good production and, when relatedness is sufficiently high, public goods benefit preferentially genotypes with high transfer ability. Both our simulations and experiments indicate that transfer is not required to occur among kin, provided that public goods still benefit kin. Public good gene mobility thus aligns the interests of chromosomes and MGEs concerning transfer, promoting gene exchange among bacteria.

## Introduction

Plasmids and other mobile genetic elements (MGEs) often bear genes involved in social interactions between cells, where the behaviour of an individual cell will impact the fitness of its neighbours (1). Some well-known examples are B-lactamases that degrade antibiotics extracellularly (2), and secreted toxins involved in bacterial virulence (3). More generally, genes coding for secreted proteins, ones likely to participate in social interactions, are overrepresented on mobile elements (4), an observation that led to research investigating the causes and consequences of social genes enrichment in the mobile gene pool.

Many secreted molecules can be characterized as “public goods” - molecules costly to produce but readily available and beneficial to not only the producer but also to other neighbouring individuals. Such public goods are subject to specific ecological dynamics, where the public good producers are threatened by the spread of “cheaters”, neighbouring individuals that benefit from the public goods without paying production costs (5). Theory and experiments have shown that horizontal transfer can counteract the invasion of non-producer cells in two ways: first, transfer of public good genes towards previously non-producing cells enforces cooperation (6) (7); secondly, in a structured population horizontal transfer promotes cooperation by increasing relatedness for mobile alleles: new producer cells arising from transfer are preferentially associated to other producers, favouring cooperation (5) (7). These processes can explain the enrichment of public good genes on mobile elements, as transfer increases public good gene fitness. They further suggest that transfer itself provides benefits at the population level by increasing total public good production, which could lead to indirect selection for horizontal transfer. However, genes on chromosomes and plasmids will experience different selective pressures, which might lead to conflicts over optimal rates of public good production (8) or transfer.

Both plasmid and host factors are involved in the regulation of transfer (9), and transfer rates are subject to multiple selective pressures. MGEs gain a selfish, direct advantage from transfer but high costs to the host can reduce vertical transmission (10), selecting for lower transfer rates (11). Host bacteria can modulate transfer on both donor and recipient sides. Selection on recipient ability depends on MGEs effects on fitness: when MGEs are deleterious, genotypes with defence mechanisms will be favoured (12), but beneficial MGEs promote genotypes with high recipient ability (13). Investment in donor ability is directly costly (14) (11). However, we have shown recently that genotypes investing in transfer can still spread when transfer occurs towards recipients sharing the alleles controlling transfer: preferential transfer of beneficial plasmids to kin can lead to indirect selection of plasmid donors (15). In the case of public-good producing MGEs, public goods produced because of transfer will themselves be available to neighbours, potentially modifying both host and plasmid fitness through kin selection. Here, we investigate if the indirect benefits brought by public good genes present on MGEs modify the selective pressures acting on transfer controlling genes. We hypothesize that conditions promoting public good production indirectly select for increased investment in transfer, from both plasmid and host sides.

## Selective pressures acting on chromosomal genes modulating transfer ability

### Mathematical model

Here we extend our previous model that described the selective pressures acting on non-mobile host genes modulating plasmid transfer in a patch-structured population (15). Briefly, we consider the fitness of a cell *i* in patch *j* characterized by three traits: donor ability *q_ij_*, recipient ability *r_ij_* and plasmid carriage *p_ij_* (*p_ij_*= 1 for plasmid bearing cells and 0 for plasmid-free cells). Transfer is modelled as follows: un-infected cells become infected at a rate proportional to the patch level frequency of plasmid-bearing cells *p_j_*, modulated by the mean donor ability of cells within the patch, *q_j_*, and the focal cell’s own recipient ability *r_ij_*. Donor ability brings a cost *c_q_* to plasmid-bearing cells, recipient ability has the effect *c_r_* on cell fitness. To describe the effects of public good genes on fitness, we explicitly model plasmids genes as having both a private effect *x_p_* on plasmid-containing cells, leading to the effect *x_p_ p_ij_* on fitness, and a public effect *y_p_* on all individuals in the patch leading to the effect *y_p_ p_j_* on fitness. For public good production genes, *x_p_* is negative (cost of public good production), and *y_p_* is positive (benefit of public good presence).

Based on this model, we can describe host cell fitness *W_ij_* following plasmid transfer as:

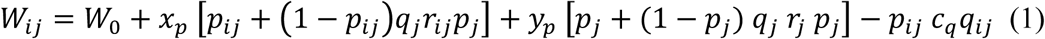

### Selection of donor ability: predictions

In this section, we consider that recipient ability has no direct effect on fitness, *c_r_ = 0*. Following our analysis in (15), we apply the Price equation to Equation (1) (see Supplementary Information for details) and obtain the regression coefficient between fitness and donor ability, *β*(*W_ij_*, *q_ij_*):

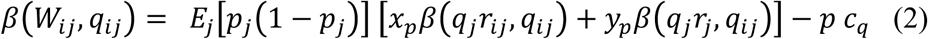

We use this equation to discuss the selection outcomes and predict when transfer will be favoured. Namely, an increase in donor ability will be favoured when *β*(*W_ij_, q_ij_*) is positive and high. Equation (2) has similarities to the result we obtained for the transfer of private good traits (15): plasmid effects on host fitness after transfer *x_p_β*(*q_j_r_ij_*, *q_ij_*)+*y_p_β*(*q_j_r_j_*, *q_ij_*), weighted by a transfer efficiency term *E_j_*[*p_j_*(1–*p_j_*)], affect indirectly the selection of donors, while transfer has the direct cost to fitness *p c_q_*. However, plasmids now act on host fitness at two levels, the individual host cell (through parameter *x_p_*, equivalent to *e_p_* in our previous model) and the group (through the new parameter *y_p_*), and we will explain their effect independently.

As previously, the effect of *x_p_* on individual cells is modulated by the factor *β*(*q_j_r_ij_, q_ij_*), a regression coefficient that describes how much transfer from donor cells is directed towards cells sharing the same donation allele, because of population structure and discrimination mechanisms. For plasmids that bring direct benefits to their host cells (*x_p_*> 0), transfer to kin favours host donor ability (15). In the case of public good genes however, we have *x_p_*< 0, and thus transfer to kin is predicted to reduce selection for donor ability.

The effect of *y_p_* describing public good benefits at the group level, is modulated by the regression coefficient *β*(*q_j_r_j_*, *q_ij_*). The factor *β*(*q_j_r_j_*, *q_ij_*) will favour cells with high donor ability *q_ij_* when they are present in patches enriched with cells with both high donor and recipient abilities, independently of the focal cell’s own recipient ability *r_ij_*. Thus, public good benefits favour donor cells when transfer within their patches is high, leading to a high level of public good production, regardless of specificity of transfer towards a particular cell type.

When the population is structured, i.e. cells with the same allele for high donor ability are clustered together within patches, donor cells will preferentially benefit from the additional public goods produced within patches after transfer. However, contrary to the case of private good genes, transfer to kin does not bring additional benefits, but additional costs to donors. Thus the population structure and specificity in transfer can also have an opposite effect on fitness, as transfer towards cells sharing high donor ability alleles will burden them with the additional cost of public good production.

Our model thus suggests that donor ability would be selected most strongly with a combination of high population structure (allowing donor cells to reap the benefits of public good production) and mechanisms ensuring that transfer occurs to other genotypes and not to their kind (avoiding the added costs of public good production). We go on to test these predictions using experiments and mathematical simulations.

### Selection of donor ability: experiments

Our model predicts that sufficient public good benefits can allow donor ability to be selected, if relatedness is high enough for benefits to favour donors and overcome the direct costs of donor ability. We now test that prediction experimentally with a synthetic system based on the one already constructed for study of interactions between transfer and public good production (7) (15). To focus on the effects of transfer on donors, we design an experiment where donor ability is directed only to unrelated recipients. Indeed, additional costs of public good production would be difficult to distinguish experimentally from the direct cost of transfer. We analyse the case of transfer to kin later, in simulations where we can distinguish costs of public goods and transfer. In our setup, several distinct strains can carry plasmids: primary plasmid-bearers (D^+^ and D^−^, differing in donor ability, neither able to receive plasmids) and secondary plasmid-bearers (recipients, R, can receive plasmids but not donate them, see Methods for more details). We compare experiments that differ in the presence of public good genes on plasmids. In the first case, D^+^ and D^−^ both bear a public-good production plasmid (P^+^), which has C_4_-HSL production genes on it, while in the second they both bear an empty control plasmid (P^−^). D^+^ can transfer the plasmid to recipients (R, leading to R_P_), D^−^ and R cells cannot (Figure S1A). The metapopulation consists of two populations differing in their D^+^ vs D^−^ ratio (Figure 1A), ensuring relatedness is positive at the metapopulation level. To start with, cells are mixed (t_0_), then transfer takes place within populations. After transfer (t_1_), populations are diluted in the presence of the antibiotic chloramphenicol (Cm), where population growth is a function of the concentration of the public good C_4_-HSL. Finally, at t_2_ equal proportions of the two populations are mixed to evaluate frequencies at the metapopulation level.

**Figure 1:**
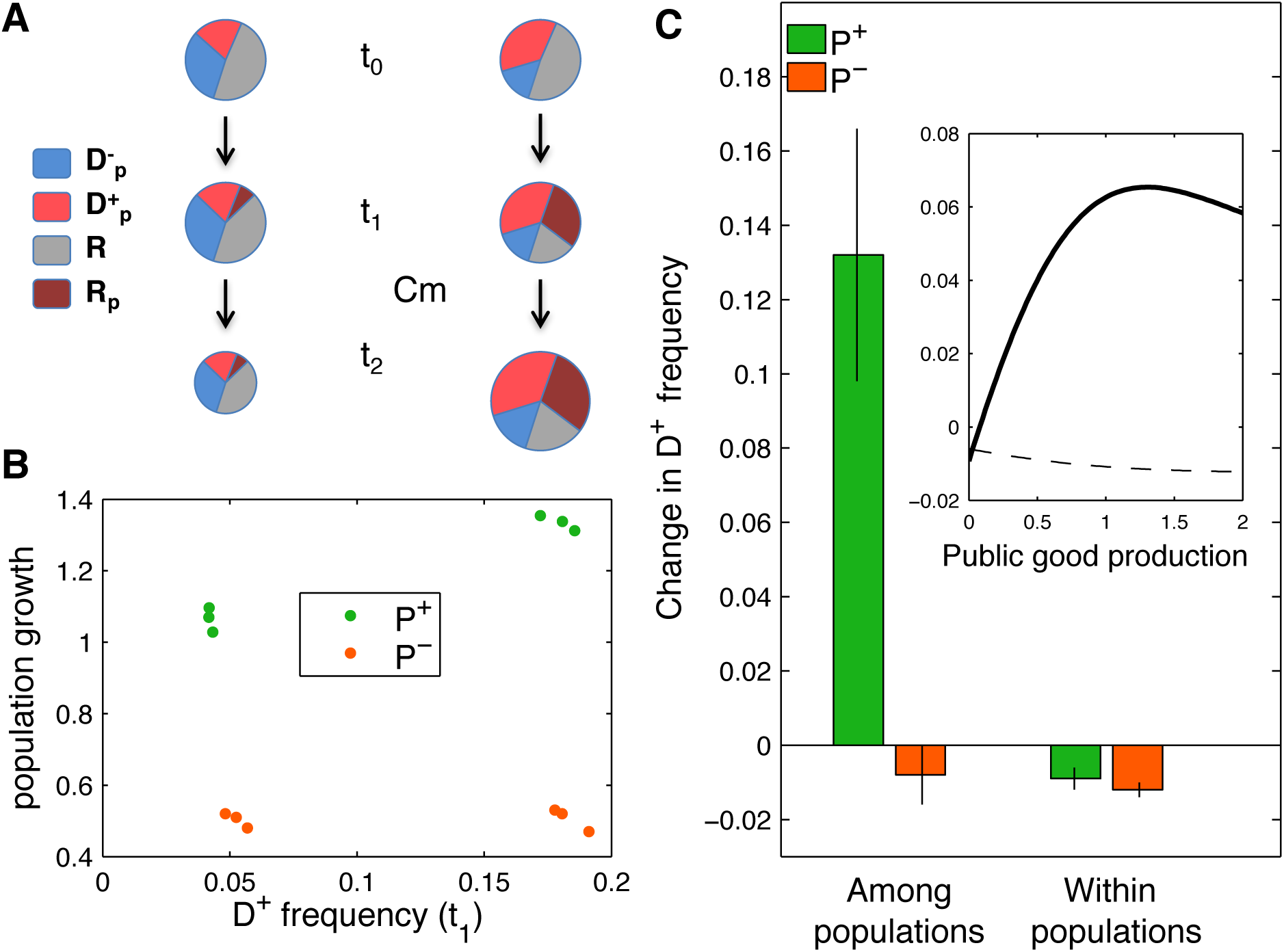
Selection of donor ability for public good genes. **A: Experimental populations.** D^+^ and D^−^ genotypes are competed, they both bear plasmids (P) that can either produce (P^+^) or not produce (P^−^) public goods (respectively green and orange in B and C panels). At t_0_, the metapopulation consists in two populations differing in their D^+^ vs D^−^ ratio, with the same proportion of recipients R. The two populations have initial D^+^/D^−^ ratios of 1/4 and 4/1 respectively, and 80% R. After growth and plasmid transfer towards R cells yielding R_P_ (t_1_), cells are confronted to chloramphenicol (Cm), and subsequent growth is dependent on previous public good production. Populations are then pooled and measured at t_2_. **B: Population growth** in each population in the presence of Cm, measured as the optical density OD_600_ after 14h of growth, is shown as a function of D^+^ proportion at t_1_. Results shown are three replicate experiments. **C: Selection of D^+^ strain.** The change in frequency of D^+^ from t_1_ to t_2_, D^+^/ (D^+^ + D^−^), is computed for strains carrying a plasmid with (green) or without (orange) public good production, and shown as mean ± s.e.m. (N=9). The inset shows the simulated change in D^+^ frequency as a function of public good production rate by plasmid genes, among (solid line) and within (dashed line) populations. D^+^ donor ability was set at *q* = 0.4 and D^−^ donor ability at *q* = 0.

Faster growth is observed in the presence of Cm for populations enriched in D^+^ cells when the transferred plasmid carries public-good genes (P^+^ plasmid) but not when it lacks them (P^−^ plasmid) (Figure 1B). The difference demonstrates that transfer increases public good production and yields benefits at the group level.

We then analyse the dynamics of the competition. Simulations (Figure 1C inset) predict that D^+^ cells are outcompeted when a non-producing plasmid is transferred, and outcompete D^−^ cells when production of public goods from the plasmid genes increases. We observe the same behaviour with our experimental system: at the metapopulation scale (among populations, Figure 1C left), plasmid donor proportion tends to decrease when donors transfer P^−^ plasmid (1% decrease, two-sided t-test for difference from 0, p = 0.32), but donors outcompete non-donor cells when they transfer P^+^ plasmid (13% increase in the frequency of D^+^, Wilcoxon signed-rank test for difference from 0, p = 0.02). Donor ability is thus selected because of public good gene transfer. To confirm that this effect is the result of metapopulation dynamics, we analyse strain dynamics at the population level. The frequency of D^+^ compared to D^−^ within each population decreases for all values of public good production in simulations (Figure 1C inset, dashed line) and experiments (Figure 1C right, 1% decrease for both P^+^ and P^−^, p = 0.014 and p = 0.008 respectively), indicating that within populations, transfer is costly. The decrease also confirms that the selection of donor ability is not a direct benefit of plasmid transfer for the donor cells within populations. Moreover, the difference between among-population and within population changes is significant for P^+^ plasmids (Wilcoxon rank-sum test, p = 0.004). Host genes promoting plasmid transfer are selected specifically when the transferred plasmid carries public good production genes and positive relatedness ensures public goods benefit donors preferentially. Selection is due to faster growth of populations enriched in plasmid donors, which more than compensates for the direct cost of transfer to donor cells.

### Selection of recipient ability: predictions

We consider here the same mathematical model as in the previous section but add a recipient ability that has a direct effect c_r_ on the host. The effect may be negative (cost of receiving) or positive (not bearing a cost of immunity mechanisms against plasmids) leading to an effect of transfer on host fitness equal to *c_r_r_ij_*. Additionally, as we focus on recipient ability, here we assume there is no cost of donor ability. With these assumptions, the regression coefficient between fitness and donor ability, *β*(*W_ij_, r_ij_*), is calculated as:

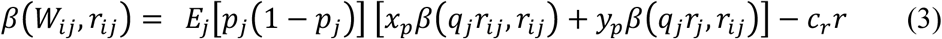

The structure of equation (3) is similar to equation (2), with two selective forces acting on recipient ability: indirect effects describing plasmid effects on fitness after transfer [*x_p_β*(*q_j_r_ij_*, *r_ij_*)+*y_p_β*(*q_j_r_j_*, *r_ij_*)],weighted by transfer efficiency *E_j_*[*p_j_*(1–*p_j_*)], and direct effects of recipient ability *c_r_r*.

Assuming that *q_j_* is independent from *r_ij_*, the term *x_p_β*(*q_j_r_ij_*, *r_ij_*)can be simplified to *q_j_x_p_*. For genes with private effects (*y_p_*=0,*x_p_* > 0), equation (3) predicts that recipient ability is selected when their benefits weighted by transfer efficiency overcome recipient ability costs, in absence of any population structuring: *E_j_*[*p_j_*(1–*p_j_*)]*q_j_x_p_* > *c_r_r*. Contrary to donor ability, under these conditions recipient ability can be directly selected for. Similar conclusions were reached in (13), where evasion of CRISPR immunity was observed when simply selecting for recipients of a beneficial plasmid.

In the case of public good genes, the term *q_j_x_p_* translates into selection against recipient ability, as receiving plasmids leads to added costs of public good production. On the contrary, *y_p_*> 0 favors recipient ability, provided that *β*(*q_j_r_j_,r_ij_*)is positive and high: increased public good production following transfer in patches enriched in recipient cells indirectly selects for recipient ability.

Equation (3) thus predicts that recipient ability for public good genes can be selected, provided that benefits and relatedness are strong enough to overcome the potential costs of recipient ability itself, and the added costs of public good production borne by recipient cells.

### Selection of recipient ability: experiments

To test our model predictions experimentally, we compete here two strains: R^+^ cells that can receive efficiently plasmids and R^−^ cells that receive them at a much lower rate, using a distinct strain as the initial plasmid donor, D_P_^+^ (in order to exclude differences in donor ability as reasons for selection, Figure S1B, see details in Methods). In simulations, we assume that R^−^ cells do not receive plasmids at all. We also assume that recipient ability confers no direct fitness cost to the host (*c_r_* = 0), but has a cost associated to plasmid burden.

The experimental setup mirrors the one presented for Figure 1, with a metapopulation consisting of two populations differing in their R^+^ vs R^−^ ratio (Figure 2A), and the same public good system providing faster growth when confronted to Cm.

**Figure 2:**
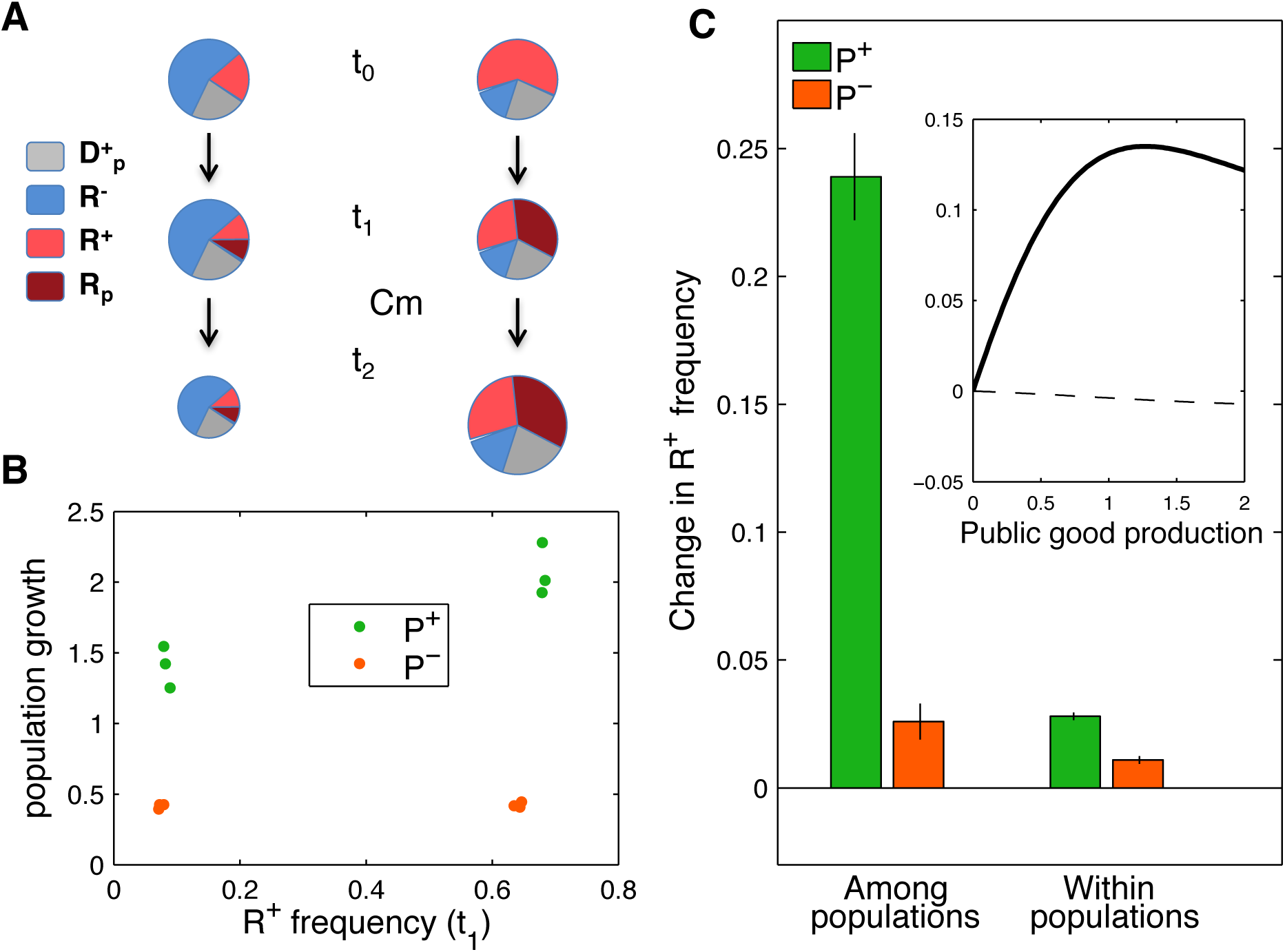
Selection of recipient ability for public good genes. **A: Experimental populations.** R^+^ and R^−^ strains are competed in a metapopulation setup. The metapopulation consists of two populations initially differing in R^+^ vs R^−^ ratio, with the same proportion of donor D^+^_P_ cells that transfer P plasmids (t_0_). Plasmids can either produce (P^+^) or not produce (P^−^) public goods (respectively green and orange in B and C panels). The two populations have initial R^+^/R^−^ ratios of 7/1 and 1/7 respectively, and 20% D^+^_P_ cells. After growth and transfer (t_1_), cells are confronted with chloramphenicol (Cm), and their subsequent growth depends on the previous public good production. Populations are then pooled and measured at t_2_. **B: population growth** is shown in each population in the presence of Cm, measured as the optical density OD_600_ after 14h of growth, as a function of R^+^ proportion at t_1_. Results shown are three replicate experiments done the same day. **C: Selection of R^+^ strain:** The change from t_1_ to t_2_ in frequency of R^+^ compared to R^−^, R^+^/ (R^+^ + R^−^), is computed for strains carrying a plasmid with (green) or without (orange) public good production, and shown as means ± s.e.m. (N = 6). The inset shows the simulated change in R^+^ frequency as a function of public good production rate of the plasmid transferred by donor cells, among (solid line) and within (dashed line) populations. R^+^ recipient ability was set at *r* = 0.4.

Differences in recipient ability effectively lead to a strong difference in the proportion of cells that receive the plasmid: in the presence of 20% donor cells, at t_1_, almost no (< 2% of the total population) R^−^ cells receive the plasmids, whereas 36% of R^+^ cells receive it. This leads to faster growth in the presence of Cm for populations enriched in R^+^ cells when the transferred plasmid carries public-good genes (P^+^ plasmid) but not when it lacks them (P^−^ plasmid) (Figure 2B).

We next analyse the dynamics of the frequency of R^+^ compared to R^−^ cells in experiments and simulations, under the same 20% initial frequency of D_P_^+^ donor cells. Simulations (Figure 2C inset) predict that R^+^ cells outcompete R^−^ cells when production of public goods from the plasmid increases. In the absence of public good production, R^+^ and R^−^ cells do equally well (as we assumed no direct cost for recipient ability, and no plasmid cost). In experiments, P^+^ plasmid recipients strongly outcompete non-recipient cells at the metapopulation scale (Figure 2C left, 24% increase in R^+^ frequency, two-sided t-test for difference from 0, p < 0.001). P^−^ recipients also slightly outcompete non-recipients (3% increase in R^+^ frequency, two-sided t-test for difference from 0, p = 0.007). Still, the difference between P^+^ and P^−^ transfer is highly significant (two-sided t-test, p < 0.001).

In simulations, R^+^ frequency decreases with the increase in the public good production within populations: recipient ability for a public good producing plasmid is costly, confirming that the selection of recipient ability is due to among-population selection. In our experiments however, there is no decline but a slight increase in R^+^ frequency within populations for both types of plasmids received: the cost of public good production added on recipients appears to be negligible. Moreover, R^+^ and R^−^ cells differ in both the F_H_ plasmid (that R^−^ bear to reduce recipient ability) and fluorescence markers they express: the net cost on fitness appears to be higher for R^−^ cells. Still, we can conclude that most of the increase in R^+^ frequency happens because of among population differences when the transferred plasmid bears public-good production genes: among- and within-populations changes are significantly different when P^+^ plasmid is transferred (two-sided t-test, p < 0.001), not when P^−^ plasmid is transferred (two-sided t-test, p = 0.08). Host genes promoting the cell recipient ability can thus be selected for when the transferred plasmid carries public-good production genes and relatedness is high enough, because of faster growth of populations enriched in plasmid recipients.

## Effect of transfer efficiency and specificity on donor selection

Next, we investigate in simulations the effect on donor selection of two parameters affecting transfer: the abundance of plasmid-bearing cells, and transfer bias towards specific cell types. To do so, we again model two strains, D^+^ and D^−^, with only D^+^ being able to transfer plasmids. Instead of considering a separate recipient strain R, D_P_^+^ cells can now transfer plasmids to either D^+^ or D^−^ cells.

First, we explore the effect of varying the initial proportion of plasmid-bearing cells, in the absence of bias in transfer from D^+^_P_ (transfer happens equally towards both D^+^ and D^−^, with a probability equal to their relative frequencies in the local population). Equation (2) predicts that selection will peak when transfer is the most efficient, at the intermediate frequency of 50% plasmid-bearing cells. With standard parameters measured from our experiment (Figure 1), selection of donors is indeed higher at intermediate frequencies (Figure 3A, D^+^ favoured when plasmid-bearing cells represent initially 0.5% to 61% of the population). Both high and low initial plasmid proportions lead to counter-selection of donors: donor ability brings no sufficient benefits in the absence of efficient transfer to overcome costs. However, diverging from equation (2) predictions, maximal selection is observed for an initial plasmid proportion of 16%.

**Figure 3:**
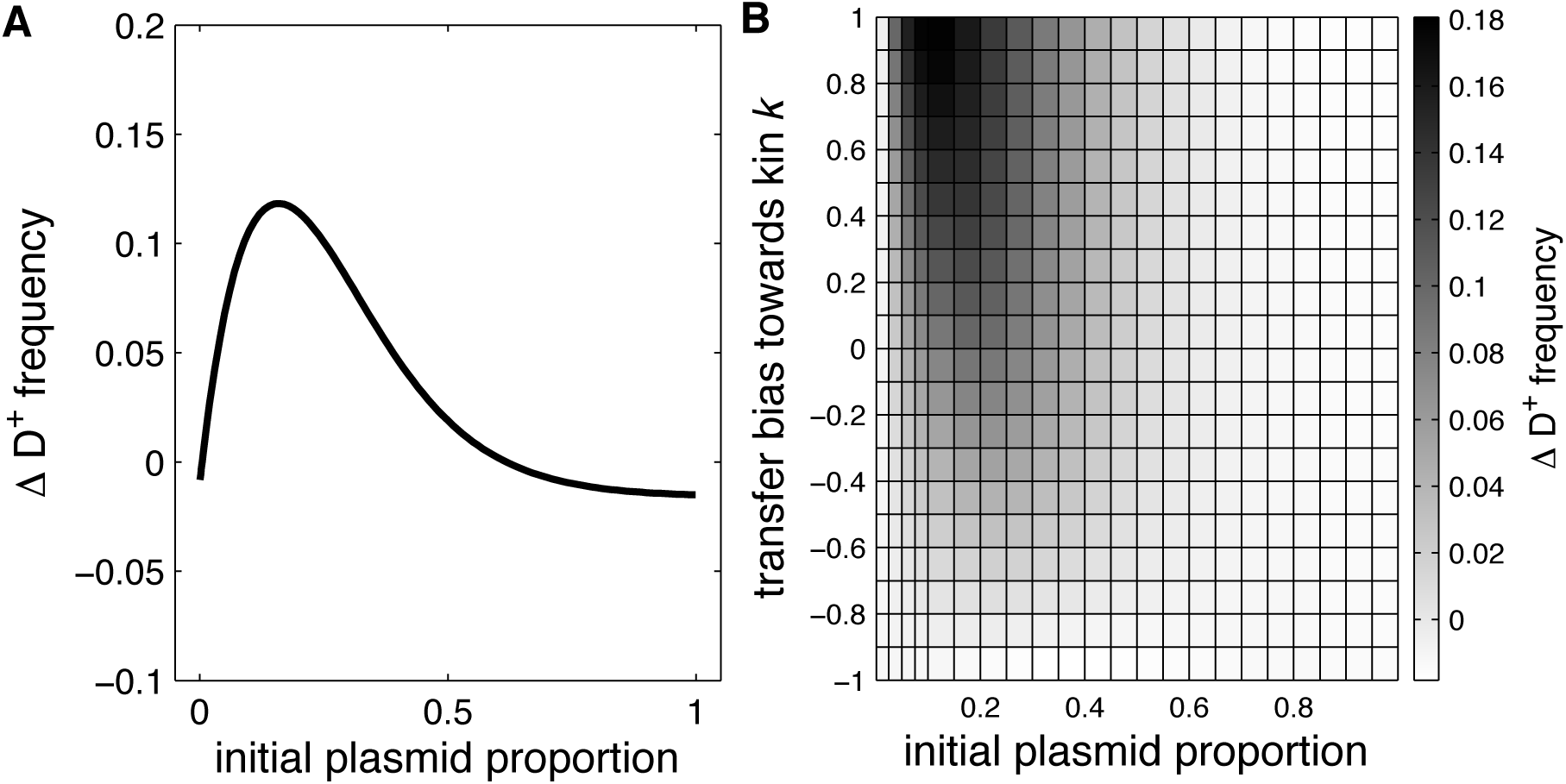
Effect of initial plasmid proportion and transfer bias on the selection for donors. In **A**, the change in donor frequency from t_0_ to t_2_ is shown as a function of initial plasmid proportion in both D^+^ and D^−^ strains, in the absence of transfer bias. In **B**, the effect of varying transfer specificity from only to D^−^ (*k* = −1) to only to D^+^ (*k* = 1) on the change in donor frequency is shown as a grayscale gradient.

To understand this divergence from theoretical expectations, we measure the efficiency of transfer (Fig S2A): the increase in plasmid-bearing cell frequency is maximal below 50%. By contrast, simulations performed with a separate, non-transferring strain as a recipient do have maximal transfer at precisely 50% initial frequency (Figure S2A). When some of the recipients can transfer back plasmids, secondary donors appear over the course of growth, increasing total transfer but also shifting the initial plasmid frequencies that maximise transfer towards lower values. Secondary transfer, not modelled in Equation (2), thus increases selection for donors at low plasmid frequencies. Moreover, selection for donor ability is based on donor-rich populations benefiting more from public goods that other populations. The relative part of public good production due to transfer is higher when initial plasmid-bearers are present in lower proportion (Figure S2B), leading to relative benefits of transferring the public good genes being maximized at plasmid-bearers proportion lower than 50%.

To investigate the effect of transfer bias in donor selection, we next introduce in the model a parameter *k*, the probability of transfer being directed preferentially towards kin (D^+^_P_ towards D^+^) vs. towards non-kin (D^+^_P_ towards D^−^) (Figure 3B). In strong contrast to Equation (2) predictions, transfer bias towards non-kin decreases selection for donor ability and transfer bias towards kin increases it. Moreover, the effect of transfer bias depends on the initial proportion of plasmid-bearing cells: selection of donors peaks at a lower plasmid proportion when transfer is biased towards kin. Again, these results can be understood by considering the phenomenon of secondary transfer, not taken into account in Equation (2). Biased transfer towards or away from D^+^ cells modifies the total transfer efficiency: D^+^ recipients behave as secondary donors, leading to more transfer especially at lower frequencies (Figure S3).

Our simulations thus uncover and underline the importance of secondary transfer for the selection of donors, a parameter not taken into account by our initial model. Selection of donor ability is the strongest when transfer is more effective generally, but also when transfer is amplified specifically among transfer-competent cells because donors transfer plasmids to their kind.

## Effect of public good genes on the selection of plasmid transfer genes

After investigating how carriage of public good genes modifies the selection of host transfer genes, we now look for the effect of public good genes carriage on plasmid transfer genes themselves.

First, we perform competitions in experiments with a reference null plasmid P^−^T^−^ that neither transfers nor produces public goods, against one of 3 plasmids with (T^+^) or without (T^−^) a transfer ability allele *oriT*, and with (P^+^) or without (P^−^) the public good allele *rhlI* (Figure S1C). Similar to previous experiments, cells first undergo a transfer phase before being confronted by the Cm antibiotic during the cooperation phase, where public good production allows faster growth in producer-rich populations. After the transfer phase (Figure 4A, left), the two plasmids with transfer allele T^+^ strongly increase in frequency compared to the null plasmid, whereas the non-transferring producer plasmid P^+^T^−^ declines in frequency (4% decrease, two-sided one sample t-test against 0, p = 0.007). P^+^T^+^ increases slightly less than P^−^T^+^ (33% vs 35%, two-sided t-test, p=0.027), likely due to the cost of public good production to P^+^T^+^ bearing cells. When public goods become beneficial during the cooperation phase (Figure 4A, right), P^+^T^+^ becomes the fittest plasmid, increasing in frequency by 46% (compared to 40% for P^−^T^+^, two-sided t-test p = 0.001). Despite public good benefits, P^+^T^−^ still declines in frequency (but not significantly, −2.5%, two-sided t-test p = 0.17). To investigate if P^+^T^+^ high fitness is linked to public good production or to direct fitness benefits within populations, we measure within-population dynamics (Figure S4). We find that the fitness patterns are similar to metapopulation ones at t_1_ (P^−^T^+^ is the fittest) but differ at t_2_, where P^+^T^+^ is less fit than P^−^T^+^. We thus conclude that the increase in P^+^T^+^ vs P^−^ T^+^ at t_2_ is a metapopulation phenomenon and requires positive relatedness. Our experiment demonstrates that carrying public good production genes can benefit a conjugative plasmid, as the fittest plasmid bears both public good and transfer alleles.

**Figure 4:**
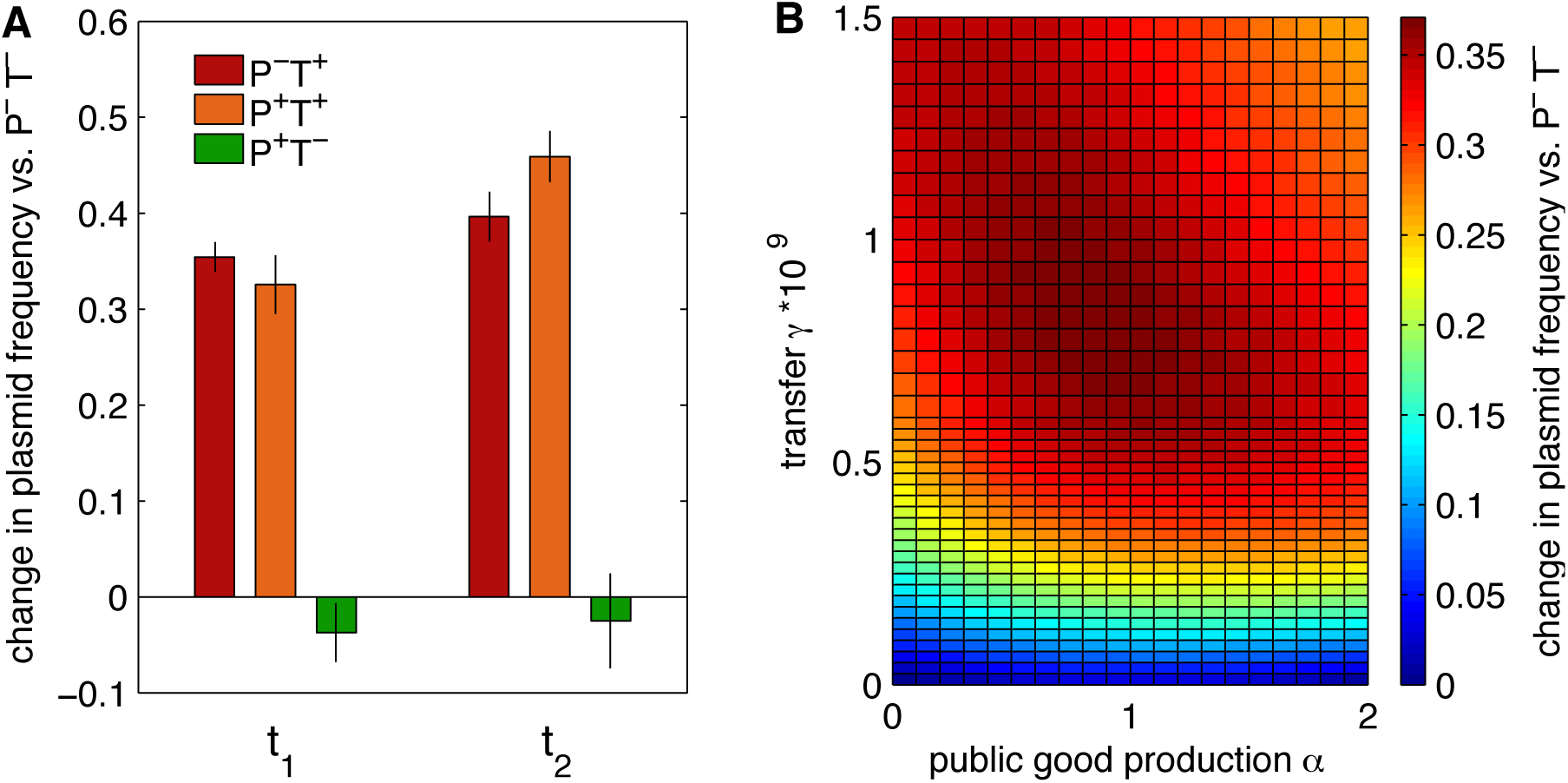
Plasmid fitness with varying transfer and public good production rates. The change of plasmid frequency is measured in comparison to a reference non-producing and non-transferring plasmid P-T^−^. 25% of cells initially bear plasmids. In **A**, the frequency of plasmids bearing or not bearing the production allele *rhlI* (P^+^ vs P^−^) and the transfer allele *oriT* (T^+^ vs T^−^) were measured experimentally, at the end of the transfer phase (t_1_) then cooperation phase (t_2_). In **B**, public good production rate α and transfer rate γ for P^+^T^+^ were varied in simulations, and P^+^T^+^ change from t_0_ is shown as a colorscale after both transfer and public good production steps (t_2_).

To analyse this result further, we measure in simulations the change in P^+^T^+^ frequency as a function of its public good production α and transfer rate γ (Figure 4B). In the absence of public good production (α = 0), increasing γ strongly favours P^+^T^+^, as the population gets progressively invaded by mobile plasmids. The effect saturates at a transfer rate around 10^−9^ mL. cell^−1^h^−1^. For non-mobile plasmids (γ = 0), P^+^T^+^ slightly benefits from intermediate production rates (α around 1). When P^+^T^+^ plasmid both transfers and produces public goods (α > 0 and γ > 0), we observe an interaction between transfer and public good production: selection of P^+^T^+^ plasmid is maximized for intermediates rates of both transfer and public good production, confirming our experimental results. Transfer and public good production act in synergy on plasmid fitness: the increase in P^+^T^+^ frequency with given transfer and production rates is greater than the sum of their independent effects. This synergistic effect happens only for intermediate rates: for higher α and γ the selection of P^+^T^+^ declines. Strikingly, for production rates α > 0, the transfer rate maximizing plasmid fitness is not the maximal transfer rate tested, contrary to what happens in the absence of public good production (α = 0). Reversely, for a given transfer rate the highest production rates also decrease plasmid fitness.

Within populations, increasing public good production lowers the frequency of P^+^T^+^ plasmid, even in the presence of transfer (Figure S5A). Public good production only brings a cost to host cells, decreasing plasmid vertical transmission, and maximal invasion happens for the highest transfer rate and no public good production, thus, the observed interaction at metapopulation scale does not arise through within-population dynamics. Among populations however, a strong interaction between transfer and public good production is apparent (Figure S5B), based on differential growth of populations, with the difference peaking for high public good production and intermediate transfer rates (Figure S5C). The effect of transfer rate can be understood by analysing plasmid spread (Figure S5D): in the s_2_ population, which is initially enriched in P^+^T^+^ plasmid (red line), plasmid spread stops increasing for high transfer rates, whereas in the s_1_ population where P^+^T^+^ is initially infrequent, plasmid invasion still increases for higher transfer rates. As a result, the difference in public good production among populations decreases with increasing transfer rate, differences among populations are thus maximized for intermediate transfer rates.

Public good production can here modulate the optimal transfer rate for a given plasmid. We now investigate the generality of this result, including the surprising fact that low transfer rates can be favoured because of among-population dynamics. In the previous scenario, we have not considered actual competition over transfer between plasmids, as P^+^T^+^ was the only mobile plasmid. We consider an additional case, where P^+^T^+^ plasmid now competes against a non-producing but highly mobile plasmid, P^−^T^high^ (Figure S6). For low public good production rates (α ≈ 0) P^+^T^+^ is outcompeted when its transfer rate is lower than the one of the competing plasmid, and outcompetes P^−^T^high^ with higher transfer rate, as expected (Figure S6A). Public good production modifies the competition, increasing P^+^T^+^ frequency for a given transfer rate. The increase is again due to among-population dynamics (Figure S6B): high P^+^T^+^ transfer rates amplify among-population variations in P^+^T^+^ frequency. However, here decreasing transfer is not favoured. Indeed, the competing plasmid also spreads within populations and public good production does not saturate with increased P^+^T^+^ transfer. Overall, public good production still benefits the mobile P^+^T^+^ plasmid in the presence of a competing highly mobile plasmid, but P^+^T^+^ optimal transfer rate is modified as the range of parameters where among-population selection is the strongest is altered.

### Effect of plasmid-free cell abundance on plasmid transfer selection

We next analyse the influence of the initial abundance of plasmid-free cells, focusing on a case where the producing plasmid P^+^T^+^ competes against P^−^T^−^ plasmid (Figure 5). When P^+^T^+^ transfer rate is low (γ ≈ 0), selection for producers increases with increasing plasmid proportion, as relatedness among public good producers gets higher. With increasing proportion of plasmid-free cells, increasingly high transfer rates maximise fitness. Within populations, P^+^T^+^ plasmid spread is highest for high transfer rates and low initial plasmid proportion (Figure S7A), because these conditions allow a complete invasion of populations by horizontal transfer. The differential success of P^+^T^+^ among populations, however, is highest when plasmids are more abundant (Figure S7B), and with intermediate transfer rates: for a given plasmid abundance, increasing transfer first increases among-population selection, however the among population change in frequency then decreases for higher values of transfer rate. The dynamics here is similar to the one described in Figure S5D: the difference in transfer efficiency in populations differing in initial plasmid proportion actually decreases as populations get completely invaded by P^+^T^+^ plasmid. The within-population component of plasmid success still increases whilst the among-population component decreases.

**Figure 5:**
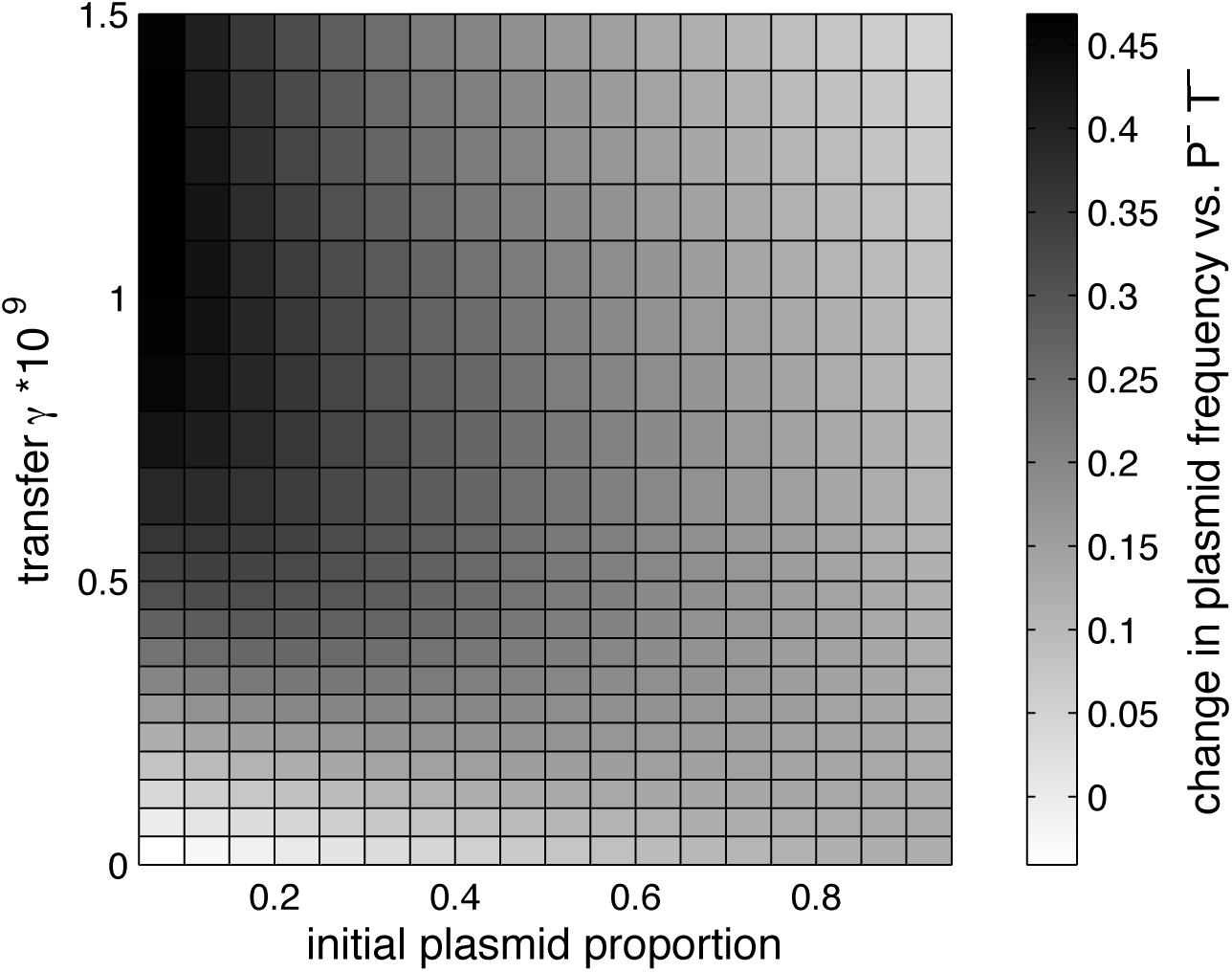
Effect of initial plasmid proportion on the selection of public-good producing plasmids. The change in frequency of P^+^T^+^, here with a public good production rate α = 1, shown as a gray scale gradient, is computed in simulations as a function of its transfer rate γ and total proportion of plasmid-bearing cells at t_0_.

The influence of public good production on plasmid fitness is thus globally dependant on the opportunities for transfer. When plasmids are initially rare, the effect of infectious transfer is predominant, and strongly increases plasmid frequency. When plasmids are more abundant, the effect of transfer on among-population dynamics becomes a significant part of plasmid selection. Synergy between plasmid transfer and production rates happens mainly when the benefits of public goods are strong enough relatively to plasmid spread to influence significantly plasmid dynamics.

## Comparison of the selective pressures acting on plasmids and chromosomes

Our separate analyses of selective pressures acting on plasmids and chromosomes show that carriage of public goods genes on plasmids can promote selection for increased transfer on both sides, providing relatedness is sufficiently high. These results imply that certain conditions align the interests of plasmids and chromosomes concerning plasmid transfer rates, when public good genes are present. To illustrate that more clearly, in the final set of experiments we perform parallel simulations with the exact same parameters, except that the alleles determining variation in plasmid transfer rate are located on either the chromosome or the plasmid: in the first case, two hosts D^+^ and D^−^ are competed; in the second two plasmids T^+^ and T^−^ are competed. T^+^ plasmids and plasmids hosted by D^+^ have the same transfer rate. Moreover, for a strict comparison with the chromosomal case, we modify the plasmid competition design relative to Fig. 4 and assume that both T^+^ and T^−^ are public good producers. We measure the effect of two key parameters on the selection of transfer (Fig 6): the public good production rate α by plasmid-located genes (x-axis); and population structure that controls the relatedness at the transfer locus (y-axis).

**Figure 6:**
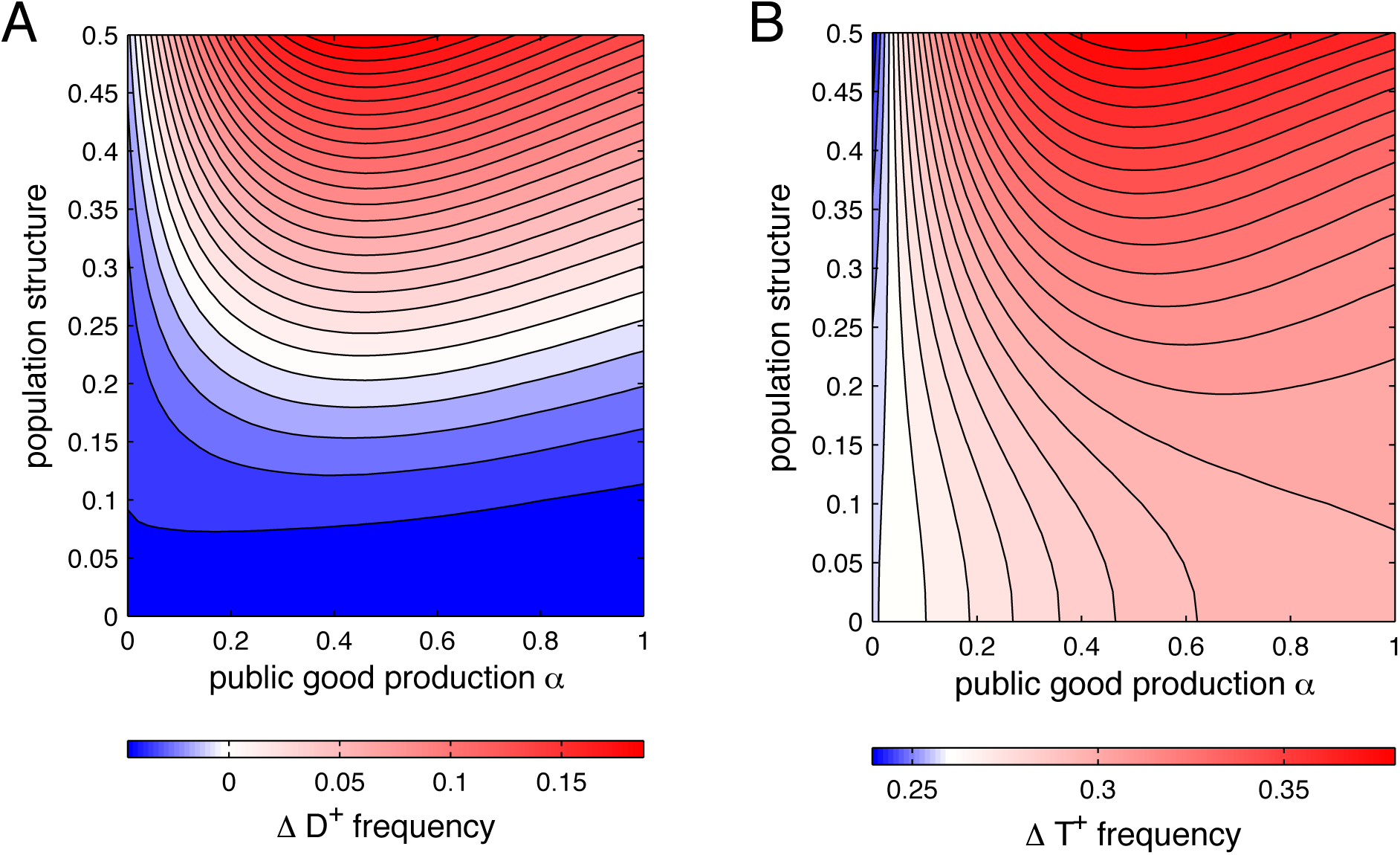
Public good genes on plasmids resolve conflicts about transfer. A genotype with transfer ability is competed with a non-transferring genotype, comparing transfer genes located on the chromosome (A) and on plasmids (B). Changes in frequency of the transferring genotype from t_0_ to t_2_ are shown as a colorscale (note that the scale is different for A and B), as a function of population structure and the level of public good production. The strength of population structure is defined as the initial difference in transfer allele frequency *f* between the two subpopulations s_1_ and s_2_: *f(D^+^)s_2_ – f(D^+^)s_1_* in A, *f(T^+^)s_2_ – f(T^+^)s_1_* in B.

When plasmids do not bear public good genes (α = 0), transfer benefits plasmids but harms chromosomal genes, showing a conflict of interests. When plasmids bear public good genes in a well-mixed population (α > 0, r = 0), the effect on fitness remains negative. Interestingly, plasmid fitness increases with increased public good production, despite the absence of population structure. That is due to the additional horizontal spread allowed by the higher population sizes achieved with public goods production (Fig S8). With increasing population structure, plasmid fitness slightly decreases, as the opportunities for transfer get more limited. Finally, with sufficient population structure, public goods increase not only T^+^ plasmid fitness through indirect benefits from transfer, as was already seen in Fig 4, but also D^+^ donor fitness. Thus, enough structure and public good production make transfer beneficial for both chromosomal and plasmid genes, aligning their interests.

## DISCUSSION

Using simulations and experiments we have shown that MGEs transfer can be selected via the benefits provided by the public good genes MGEs encode. This indirect selection acts on all loci involved in modulating transfer rates: non-mobile loci from the host on both donor and recipient sides, and plasmid genes themselves.

For host genes, enhancing transfer of public good genes is an altruistic behaviour with no direct benefits (Fig 1, Fig 2), and the condition for its selection can be expressed in a form analogous to Hamilton’s rule (16): the benefits of public goods produced by neighbours, weighted by the efficiency of public good gene transfer and relatedness, have to outweigh the costs of transfer to the host. Contrary to the case of private good genes (15), selection does not require that transfer happens towards related cells as a cell can get the benefits from altruism without itself bearing the public good plasmids. Nevertheless, relatedness must still be high in the population: cells sharing alleles that favour transfer have to be associated more than randomly, in order for benefits of additional public goods produced following transfer (to either related or unrelated cells) to feedback preferentially on them. This is comparable to the conditions required for interspecies mutualism, where cooperation towards another species necessitates within-species relatedness, so that the response from the other species benefits preferentially cells related to the initial cooperators (17) (18).

At first sight, transfer to kin is deleterious, leading to increased costs of public good production. But other effects can modify that situation. For instance, our simulations underline another effect of transfer to kin by efficient donors: secondary plasmid-bearers themselves transfer efficiently and enhance public good production compared to a scenario where initial donors transfer only to non-kin (Figure 3B). Alternatively, transfer might be increased from secondary donors irrespective of them being kin or non-kin because of transitory derepression of plasmid transfer in new recipients, leading to epidemic spread in new niches favouring strong public good production (19).

In our analysis, we only considered a scenario where public goods benefit every cell type in the same way. However, transfer to non-kin could also be a form of manipulation of recipient cells that do not benefit from public goods by the donor strain. The most striking example of this is T-DNA transfer from the Ti plasmid of *Agrobacterium tumefasciens* to host plant cells. T-DNA integrated in the plant genome induces tumour formation and the production of specific metabolites, used by *Agrobacterium* cells only (20). In this example, there is selection of T-DNA donor ability, but obviously not of recipient ability by the parasitized plant. In the context of transfer between competing bacteria, harmful effects of transfer and public good production on recipients could also be interpreted as spiteful transfer by the donors towards competitors. For instance, Shiga toxin genes in *Escherichia coli* are borne by temperate phages; horizontal transfer in the intestine (21) can lead to increased public good production (22) but also brings a competitive advantage to the donors as recipients are killed in the process. Recipients can thus simultaneously be competitors and be exploited for public goods production. Both mechanisms have been shown recently in a system where Shiga toxins provide defences against predators (23.) Such situation could evolve towards or from a cooperative scenario. For instance, selection for transfer could first act within species, but the mutations leading to increased public good production and damage to unrelated recipients could then be favoured in a plasmid coevolving with a given donor strain.

For plasmids genes, the indirect benefits of transfer brought by public goods add up to the direct benefits of transfer on fitness, which themselves can be influenced by several factors, including the availability of recipients, or the amount of competition among plasmids (24). Through its effects on among-population selection, transfer ability can be more strongly selected when public good production genes are present on mobile plasmids (Figure 4). However, that does not necessarily mean that plasmids will be selected towards higher transfer rates: our simulations also show that bearing public good genes can decrease a plasmid’s optimal transfer rate when that maximizes the variability in producer frequencies among populations. Thus, bearing public good genes will generally favour plasmid mobility, but can modulate the optimal transfer rate in different directions. Moreover, the relative importance of selective pressures linked to public good production will depend on the strength of the direct benefit given by infectious spread (highest when plasmid-free cells are abundant, Figure 5) and on population structure (assortment is needed for selection of public good production).

We thus show that public goods gene carriage on plasmids promotes increased transfer, on both chromosomal and plasmid sides, leading to an alignment of interests when relatedness, required for public good benefits to feedback on transfer alleles, is high enough (Figure 6).

Continued benefits of transfer for both hosts and plasmids requires that plasmids do not spread to fixation. Several mechanisms contribute to maintaining plasmid-free cells in bacterial populations, as we have discussed previously (15). Public good plasmids should be especially prone to high loss rates from populations due to competition with cheater genotypes. That regular input of plasmid-free cells will promote continued selection for transfer – both direct selection for infectious plasmids, and the indirect selection we demonstrate here in structured populations. Transfer will simultaneously favour public good gene maintenance through increasing relatedness at mobile public good loci (7). For the host lineages, not bearing public good plasmids constantly may allow reduced costs, whilst horizontal transfer would ensure that plasmids spread and lead to high public good production when needed. Plasmid prevalence and ecological dynamics are rarely well known in natural settings, but potential public good loci often appear to undergo dynamic loss and transfer. For instance, *Agrobacterium* plasmids can be lost at high rates (25). Many secreted toxins (likely to behave as public goods) are both transferable and unstable, like virulence toxins in *Escherichia coli* (26) (27) or effector genes from the plant pathogen *Pseudomonas syringae* (28).

In conclusion, we demonstrate here that the benefits of public good cooperation can promote enhanced horizontal gene transfer in bacteria. This new selective pressure will add to the direct infectious benefits of transfer for MGEs themselves (10), and to the kin selection of altruistic transfer for the host (15). Being specific to public good genes, that benefit can explain the prevalence of those genes on MGEs (4), a localisation beneficial to both hosts and plasmids. Moreover, transfer to non-kin is also selected because of increased public good production, potentially selecting for transfer among species and the evolution of plasmids with broader host range.

## Materials and methods

### Plasmids and strains

As an example of public good genes, we use the synthetic public good system characterized in previous studies (29, 30). The background strain is JC1191, an *Escherichia coli* strain that responds to the presence of the quorum-sensing auto-inducer C_4_-HSL by growing in low concentrations of chloramphenicol (Cm). An att::rhl-catLVA(Sp^R^) segment is responsible for Cm resistance through expression of the *rhlR* gene and a Rhl auto-inducer-responsive promoter (Prhl^*^) driving an unstable version of *cat*.

Plasmids used in experiments are shown in Figure S1, relevant properties are the following:

The gene *rhlI* encodes the rhlI enzyme, which catalyses the production of C_4_-HSL. We use plasmids with the *rhlI* gene as public-good plasmids (P^+^) and plasmids without *rhlI* as control plasmids (P^−^). Moreover, transferable plasmids (T^+^) carry the wild-type origin of transfer, oriT sequence of F plasmids, allowing mobilization by helper F plasmids.

To modulate transfer ability, cells bear either no F plasmid, or F_HR_ or F_HE_ helper plasmid (7). Both F_HR_ and F_HE_ are mutants of the F plasmid pOX38::Tc (31) with two point mutations in *oriT* sequence that lead to strongly reduced self-mobilization. F_HE_ has the wild-type *traS* gene encoding F plasmid entry exclusion (thus F_HE_, for helper with entry exclusion), leading to very low recipient ability of cells bearing it. F_HR_ bears a deletion of the *traS* gene: recipient cells bearing F_HR_ are able to receive plasmids efficiently (100-fold more efficiently than cells bearing F_HE_) and behave as secondary donors of oriT-bearing plasmids (thus F_HR_, for helper receiver plasmid).

Finally, plasmids and strains are identified by flow cytometry, through combinations of fluorescent markers, whose construction is described in (7) and (15). Plasmids bear GFP or YFP markers; one of two strains bears a red fluorescence marker: for experiments concerning donor ability, recipients R bear pSB3K3-RFP plasmid (7). For other experiments, one of the strains is marked with the *td*-*Cherry* gene under control of a pRNA1 promoter, inserted in the attTn7 site of the chromosome (described in (15)). Strain properties and fluorescent markers for identification by flow cytometry are summarized in Figure S1 for each competition experiment setup.

### Growth and experiment conditions

Cells were grown in Luria-Bertani (BD Difco) medium with 25 μg/mL spectinomycin (Sp, Sigma-Aldrich) and with or without 50 μg/mL kanamycin (Kn, Sigma-Aldrich) and 6.25 μg/mL chloramphenicol (Cm; Sigma-Aldrich). Experiments were conducted under well-mixed conditions with 5mL medium in 50 mL tubes (Sarstedt). For competition experiments, strains were mixed at various ratios (vol/vol) and first grown from a 10-fold dilution from stationary phase cultures (t_0_) at 35°C, up to an optical density of 3. This step was repeated once with 5-fold dilution in order to increase plasmid transfer. Cultures were then diluted 10-fold and grown overnight until stationary phase at 30°C (t_1_). Finally, cultures were diluted 100-fold at t_1_ and grown for 12 to 16 hours at 30°C until t_2_. For competition experiments between donors (Figure 1), the growth medium contained Sp and Kn from t_0_ to t_1_, and Sp, Kn, and Cm from t_1_ to t_2_. For competition experiments between recipients (Figure 2) and between plasmids (Figure 4), the growth medium contained Sp from t_0_ to t_1_, and Sp and Cm from t_1_ to t_2_.

Cultures were analysed for strain and plasmid proportions by flow cytometry, as described in (7). Data acquisition was performed on the Cochin Cytometry and Immunobiology Facility. Optical density at 600nm was measured on a BioPhotometer plus (Eppendorf). Frequencies at the metapopulation level were measured by pooling equal volumes of populations, effectively taking into account differential growth among populations. Frequencies within populations were computed as the mean of each individual frequency within populations, in order to exclude the effect of differential growth among populations.

### Simulations

Our models were designed to mimic experimental conditions of strain growth and plasmid transfer, similarly to our previous work (7, 15). Growth follows a logistic function depending on total cell density *n_tot_* and saturating at carrying capacity *K*. After a transfer phase without antibiotics (from t_0_ to t_1_), cells grow in the presence of Cm (from t_1_ to t_2_), making public goods beneficial.

Plasmid transfer happens at a basal rate γ, following a mass-action law, and saturates at carrying capacity *K*. We assume that the presence of a plasmid within a cell prevents superinfection by another plasmid, so do not consider here within-cell plasmid competition. Strains are characterized by donor ability *q* or recipient ability *r*, which modulate the effective transfer of plasmids *γ_e_*:*γ_e_* = *q*×*γ* when varying donor ability, *γ_e_* = *r*×*γ* when varying recipient ability. We assume a cost of transfer for the donor cell *c_q_* so that the cost on growth is proportional to donor ability, but no cost of receiving plasmids (*c_r_* = 0). Public good production has a cost *c_p_* to host cells, and leads to the benefit *b_p_* during the second phase of growth. All costs act on growth rate. In the presence of Cm, growth is modelled as proportional to the proportion of public-good producing cells at a given time in the local population (*p_tot_/ n_tot_*), and to the public good production rate *α*. Noting the basal growth rate ψ_P_ for plasmid-bearing cells and ψ_Ø_ for plasmid bearing cells, plasmid effects on growth are summarized below:

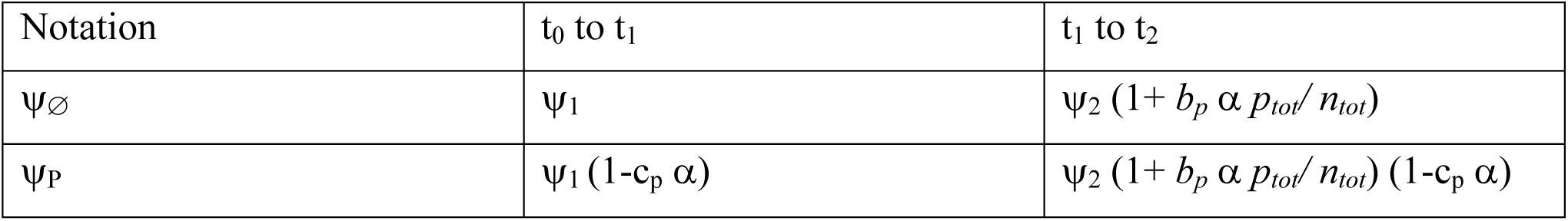

| Notation | $t_0$ to $t_1$ | $t_1$ to $t_2$ |
| --- | --- | --- |
| $\psi_\emptyset$ | $\psi_1$ | $\psi_2 (1 + b_p \alpha p_{tot}/n_{tot})$ |
| $\psi_p$ | $\psi_1 (1 - c_p \alpha)$ | $\psi_2 (1 + b_p \alpha p_{tot}/n_{tot}) (1 - c_p \alpha)$ |

Equations of plasmid and strain dynamics are given in SI Text, as well as chosen parameter values. Computer simulations were conducted using MATLAB.

## Acknowledgements

We thank all members of Institut National de la Santé et de la Recherche Médicale U1001 and Angus Buckling for advice and discussions, John Chuang for providing JC1191 strain and pZS*2R-GFP,rhlI plasmid, Didier Mazel for providing the pOX38::Tc plasmid, Chantal Lotton and the Cochin Cytometry and Immunobiology Facility for technical help.

## References

1. Rankin DJ, Rocha EPC, & Brown SP (2010) What traits are carried on mobile genetic elements, and why? Heredity 106(1):1–10.

2. Barlow M & Hall BG (2002) Phylogenetic Analysis Shows That the OXA B-Lactamase Genes Have Been on Plasmids for Millions of Years. Journal of Molecular Evolution 55(3):314–321.

3. Novick R (2003) Mobile genetic elements and bacterial toxinoses: the superantigen-encoding pathogenicity islands of *Staphylococcus aureus*. Plasmid 49(2):93–105.

4. Nogueira T, et al. (2009) Horizontal Gene Transfer of the Secretome Drives the Evolution of Bacterial Cooperation and Virulence. Current Biology 19(20):1683–1691.

5. West SA, Griffin AS, Gardner A, & Diggle SP (2006) Social evolution theory for microorganisms. Nature Reviews Microbiology 4(8):597–607.

6. Smith J (2001) The social evolution of bacterial pathogenesis. Proceedings of the Royal Society B: Biological Sciences 268(1462):61–69.

7. Dimitriu T, et al. (2014) Genetic information transfer promotes cooperation in bacteria. Proceedings of the National Academy of Sciences 111(30):11103–11108.

8. Mc Ginty SE & Rankin DJ (2012) The evolution of conflict resolution between plasmids and their bacterial hosts. Evolution 66(5):1662–1670.

9. Frost LS & Koraimann G (2010) Regulation of bacterial conjugation: balancing opportunity with adversity. Future Microbiology 5(7):1057–1071.

10. Stewart FM & Levin BR (1977) The population biology of bacterial plasmids: a priori conditions for the existence of conjugationally transmitted factors. Genetics 87(2):209–228.

11. Haft RJF, Mittler JE, & Traxler B (2009) Competition favours reduced cost of plasmids to host bacteria. The ISME journal 3(7):761–769.

12. Levin BR (2010) Nasty Viruses, Costly Plasmids, Population Dynamics, and the Conditions for Establishing and Maintaining CRISPR-Mediated Adaptive Immunity in Bacteria. PLoS Genetics 6(10):e1001171.

13. Jiang W, et al. (2013) Dealing with the Evolutionary Downside of CRISPR Immunity: Bacteria and Beneficial Plasmids. PLoS Genetics 9(9):e1003844.

14. Turner PE, Cooper VS, & Lenski RE (1998) Tradeoff Between Horizontal and Vertical Modes of Transmission in Bacterial Plasmids. Evolution 52(2):315–329.

15. Dimitriu T, et al. (2016) Indirect Fitness Benefits Enable the Spread of Host Genes Promoting Costly Transfer of Beneficial Plasmids. PLOS Biology 14(6):e1002478.

16. Hamilton WD (1964) The Genetical Evolution of Social Behaviour. I. Journal of Theoretical Biology 7:1–16.

17. Frank SA (1994) Genetics of mutualism: the evolution of altruism between species. Journal of Theoretical Biology 170(4):393–400.

18. Foster KR & Wenseleers T (2006) A general model for the evolution of mutualisms. Journal of Evolutionary Biology 19:1283–1293.

19. Dimitriu T, Misevic D, Lindner AB, & Taddei F (2015) Mobile genetic elements are involved in bacterial sociality. Mobile Genetic Elements 5(1):7–11.

20. Platt TG, Fuqua C, & Bever JD (2012) Resource and Competitive Dynamics Shape the Benefits of Public Goods Cooperation in a Plant Pathogen. Evolution 66(6):1953–1965.

21. Gamage SD, Strasser JE, Chalk CL, & Weiss AA (2003) Nonpathogenic *Escherichia coli* can contribute to the production of Shiga toxin. Infection and immunity 71(6):3107–3115.

22. Goswami K, Chen C, Xiaoli L, Eaton KA, & Dudley EG (2015) Coculture of *Escherichia coli* O157:H7 with a Nonpathogenic *E. coli* Strain Increases Toxin Production and Virulence in a Germfree Mouse Model. Infection and Immunity 83(11):4185–4193.

23. Aijaz I., Koudelka, GB (2018) Cheating, facilitation and cooperation regulate the effectiveness of phage-encoded exotoxins as antipredator molecules. MicrobiologyOpen e00636. doi:10.1002/mbo3.636.

24. Smith J (2011) Superinfection drives virulence evolution in experimental populations of bacteria and plasmids. Evolution 65(3):831–841.

25. Belanger C, Canfield ML, Moore LW, & Dion P (1995) Genetic analysis of nonpathogenic *Agrobacterium tumefaciens* mutants arising in crown gall tumors. Journal of bacteriology 177(13):3752–3757.

26. Bielaszewska M, et al. (2007) Shiga Toxin Gene Loss and Transfer In Vitro and In Vivo during Enterohemorrhagic *Escherichia coli* O26 Infection in Humans. Applied and Environmental Microbiology 73(10):3144–3150.

27. Zhang W, et al. (2013) Lability of the pAA Virulence Plasmid in *Escherichia coli* O104:H4: Implications for Virulence in Humans. PLoS ONE 8(6):e66717.

28. Neale HC, et al. (2016) A low frequency persistent reservoir of a genomic island in a pathogen population ensures island survival and improves pathogen fitness in a susceptible host. Environmental Microbiology 18(11):4144–4152.

29. Chuang JS, Rivoire O, & Leibler S (2010) Cooperation and Hamilton’s rule in a simple synthetic microbial system. Molecular Systems Biology 6.

30. Chuang JS, Rivoire O, & Leibler S (2009) Simpson’s Paradox in a Synthetic Microbial System. Science 323(5911):272–275.

31. Anthony KG, Sherburne C, Sherburne R, & Frost LS (1994) The role of the pilus in recipient cell recognition during bacterial conjugation mediated by F-like plasmids. Molecular microbiology 13(6):939–953.

